# Assimilation efficiencies and elimination rates of silver, cadmium and zinc accumulated by trophic pathway in *Gammarus fossarum*

**DOI:** 10.1101/2023.07.14.549054

**Authors:** Ophélia Gestin, Christelle Lopes, Nicolas Delorme, Laura Garnero, Olivier Geffard, Thomas Lacoue-Labarthe

## Abstract

To improve the assessment of metal toxicity in aquatic organisms, it is important to consider the different uptake pathways (i.e. trophic or aqueous). The bioaccumulation of dissolved metals such as Cd and Zn in gammarids is beginning to be well described. However, there are very few data on the contribution of the dietary pathway, and its associated toxicokinetic parameters. Among these, the assimilation efficiency (AE) is an essential parameter for the implementation of models that take the trophic pathway into account. This study aims to estimate the assimilation efficiencies and elimination rates of two types of food, i.e. alder leaves and chironomid larvae, contaminated with three metals (Ag, Cd and Zn) of major concern for the Water Framework Directive (WFD). The pulse-chase-feeding method was used. Gammarids were fed with alder leaves or chironomid larvae previously contaminated with ^110m^Ag, ^109^Cd or ^65^Zn, for a short period of time (1 to 5 hours), followed by an elimination phase of 14 days. At different time points, the gammarids were placed alive on the gamma detector to individually quantify whole body concentrations of ^110m^Ag, ^109^Cd or ^65^Zn. Our results indicate that: i) Cd has the highest assimilation efficiency (39% for leaves and 19% for larvae), followed by Zn (15% for leaves and 9% for larvae) and Ag (5% for leaves); ii) for Cd and Zn, the AE were higher when gammarids were fed with leaves than with larvae; iii) the elimination rates of metals seem to depend more on the food matrix than on the metal assimilated; and thus iv) the biological half-life calculated from the k_es_ is 5.1 days for Ag, between 4.9 and 13 days for Cd and between 3.8 and 13 days for Zn.

## 1. Introduction

In the past decades, a number of works have demonstrated the strong capacity of some freshwater organisms to accumulate contaminants from the aquatic environment (Timmermans et al., 1992; Dutton & Fisher, 2011). Consequently, some freshwater invertebrates have been used as bioindicators of good ecological status for French freshwater systems, based on the chemical concentrations recorded in the whole body (Besse et al., 2012; Pellet et al., 2014; Lebrun et al., 2020). The ubiquitous gammarid *Gammarus fossarum* is widely used for this property, as it accumulates metals as a grazer of leaf litter and is an opportunistic predator of invertebrate prey (Kunz et al., 2010; Pellet et al., 2014; Filipović Marijić et al., 2016). Although the bioaccumulation processes of dissolved metals have been extensively documented (Vellinger et al., 2012; Urien et al., 2017; Gestin et al., 2022, 2023), less is known about the dietary assimilation and elimination of metals in this species. As exposure conditions are more difficult to standardise than for aqueous exposures, the dietary route in aquatic invertebrates is often poorly understood. However, the dietary pathway could account for a significant metal entry in invertebrates (Borgmann et al., 1989; Pellet et al., 2014), which highlights the need to consider this contribution in order to avoid misinterpretation of recorded metal concentrations in this species (Franklin et al., 2005; Besse et al., 2013; Lebrun et al., 2015; Conti et al., 2016; Hadji et al., 2016).

The few studies focusing on freshwater invertebrates have shown that the uptake from the trophic pathway predominates for some metals and metalloids, such as for Cd, Cu and Se in *Hyallela azteca* (Borgmann et al., 2007) or Cd in *Gammarus pulex* (Pellet et al., 2014). In *Daphnia magna*, however, the uptake from the aqueous pathway predominates for Zn (Memmert, 1987). Previous results suggest that metal accumulation, detoxification and elimination, and thus potential toxicity, are strongly dependent on the uptake pathway. In this sense, Croteau & Luoma (2009) called for the need to delineate the bioaccumulation mechanisms and toxicokinetics in different organisms, as these processes are metal and species specific.

Assimilation efficiency (AE) is one of the most important parameters used to quantify trophic transfer. This parameter directly reflects the amount of metals available from food and is a key physiological parameter for understanding the trophic transfer mechanisms (Croteau & Luoma, 2009). The AE could be partly defined by the fraction of metal in food that crosses the intestinal epithelial barrier to accumulate in the body of the consumer (Wang & Fisher, 1999a; Nunez-Nogueira et al., 2006; Croteau & Luoma, 2008). AE values can be used to estimate the influence of consumer species, contaminants, temperature, etc. (Wang & Fisher, 1999a; Wang & Wong, 2003). Among these, the type of food can greatly influence the AE of metals and thus their bioaccumulation efficiency (Wang & Fisher, 1999a; Nunez-Nogueira et al., 2006; Dubois & Hare, 2009; Pouil et al., 2016). It has already been established that metals from plant matrices are less assimilated by organisms than elements of animal origin, because the digestion of compounds such as fibres or hemicelluloses is slower and more complex (Xu & Pascoe, 1994; Nunez-Nogueira et al., 2006). It is therefore important to consider dietary diversity in omnivorous organisms, such as gammarids, in trophic pathway studies.

The pulse-chase-feeding approach is commonly used to determine the AE values (Calow & Fletcher, 1972; Warnau et al., 1996; Metian et al., 2007; Pouil et al., 2017) and consists of exposing the organism once to a labelled food source and tracking the elimination of the tracer over time (Wang & Fisher, 1999a; Pellet et al., 2014). A kinetic modelling approach is applied to the depuration data to determine the metal-specific AE and the depuration rates (k_e_). The pulse-chase-feeding technique is widely used and adapted to work with different tools, such as isotopic tracers (stable and radioactive ones) (Wang & Fisher, 1999a; Baines & Fisher, 2002; Pouil et al., 2017). In the case of metals, gamma-emitting radiotracers are among the most powerful tracers for accurately quantifying small amounts of metal. They also allow individual tracking of metal uptake and loss over time, as whole-body gamma detection is non-destructive (Wang & Fisher, 1999a). Furthermore, it is conventionally accepted that radioactive isotopes of metals have identical properties to their stable counterparts (Warnau & Bustamante, 2007). This confers the ability to assess metal accumulation and elimination rates by measuring only the concentrations of radioisotopes, thus avoiding the measurement of the background of stable metals (Cresswell et al., 2017).

A toxicokinetic model for the aqueous pathway has already been developed for zinc and cadmium (Gestin et al., 2022). However, to be able to set up this kind of model considering the trophic pathway, it is important to have preliminary data concerning the AE of metals. In this framework, the aim of this study was to fill the knowledge gap on metal assimilation efficiency in gammarids, by determining the AE values and the elimination rates of Ag, Cd and Zn in the amphipod *G. fossarum*. A pulse-chase-feeding method was used, taking into account two contrasting types of food in the gammarid diet: alder leaves and chironomid larvae. After a short period of exposure to radiolabelled food, gammarids were transferred to depuration conditions and metal loss in tagged individuals was monitored for 14 days. Nonlinear least squares (NLS) models were fitted to the experimental data to determine the AE, k_e_ and biological half-life of metals in gammarids. The values were then compared to discuss the influence of metal and food type on dietary bioaccumulation.

## 2. Material and methods

### 2.1 Organisms and alder leaves: origins, collection and maintenance

Adult male gammarids (*Gammarus fossarum*: 27.3 ± 5.7 mg wet weight) were sampled from an uncontaminated watercress pound (Saint-Maurice-de-Rémens, France) (Gestin et al., 2021). They were then transported to the LIENSs laboratory premises (La Rochelle), where they were kept under constant bubbling in Evian^®^ water (see characteristics in Table S1) with a dark/light cycle of 8:16h, at a temperature of 12.0 ± 0.5°C. They were fed with alder (*Alnus glutinosa*) leaves *ad libitum* and kept for a 7-day acclimation period before the experiments.

Alder leaves (*A. glutinosa*) used for trophic exposure were collected from a clean site (Les Ardillats, France). They were leached (i.e. soaked in well water for several days, with daily renewal of the water) in the ecotoxicology laboratory of INRAE Lyon-Villeurbanne in order to depurate the toxic tannins for gammarid. The wet leaves were then cut into discs of 6 mm diameter, avoiding the large veins. Finally, the leaf discs were placed in a container filled with well water (see characteristics in Table S1) and transferred to the LIENSs laboratory.

Chironomid (*Chironomus riparius*) egg masses from the INRAE laboratory were transferred to the LIENSs laboratory. The eggs were placed in beakers containing powdered silica and filled with Evian^®^ water and kept at room temperature (about 20°C) for few days until hatching. The larvae were then fed with fish pellets until the third larval instar.

### 2.2 Radiotracers

Throughout the experiment, the rearing and laboratory materials were decontaminated with 1/10 diluted detergent (Decon^®^ 90 solution), 1/10 diluted HCl solution (hydrochloric acid S.G. 32%, certified AR for analysis; Fischer Scientific^®^) and distilled water. The radioisotope carrier-free solutions were purchased from Eckert & Ziegler Isotope Products Inc., Valencia, USA. The ^109^Cd and ^65^Zn solutions were in their chloride forms (i.e. CdCl_2_ and ZnCl_2_), 0.1M HCl and the ^110m^Ag solution was in its nitrate form (i.e. AgNO_3_), 0.1M HNO_3_. These three stock solutions were diluted to intermediate solutions to achieve appropriate radiotracer concentrations in the experimental setup. These radiotracer additions to the media did not change the pH of the water. For each radioisotope, the detectors were calibrated with two in-house liquid standards of the appropriate geometry*, i.e*. a 500 µL “gammarid” geometry and a 10 mL “water” geometry. Two NaI detectors coupled to InterWinner 7.0 software (ITECH Instruments^®^) were used to count the radioisotope activities (in Bq, Bq.g^-1^ or Bq.mL^-1^) present in all samples.

### 2.3 Contamination of food

The leaf discs (∼ 108 in total, i.e. 36 for each metal) were placed in 10 mL of Evian^®^ water contaminated with 2010, 2030 or 2000 Bq.mL^-1^ of ^109^Cd, ^65^Zn and ^110m^Ag, respectively, for 7 days. They were then placed in 0.1 L beakers of clean Evian^®^ water for 5 days to remove the weakly adsorbed radioisotopes on the leaf walls. At the end of the procedures, 12 discs for each metal were randomly counted by gamma spectrometry at the end of these procedures, to follow the radiolabeling of the leaf.

At the end of the third instar (i.e. L3, when they start to turn red), nine pools of 10 chironomid larvae each (i.e. 90 larvae in total) were collected, placed in 100 mL beakers filled with Evian^®^ water (i.e. n=10 per beaker) and exposed for 4 days to 201, 203 or 200 Bq.mL^-1^ for ^109^Cd, ^65^Zn and ^110m^Ag (3 pools per radiotracer). This short exposure time was chosen to avoid the emergence of radiolabelled larvae in mosquitoes in the laboratory. At the end of the exposure period, the larvae were gently rinsed (i.e. rapidly soaked in 3 successive baths of clean water), then individually frozen and all counted by gamma-ray spectrometry. No radiotracer activity was detected in chironomid larvae exposed to ^110m^Ag.

### 2.4 Experimental procedure to obtain assimilation efficiencies of gammarids

The Ag AE value and elimination rate were not determined with chironomids as food source. Prior to the pulse-chase feeding experiment, gammarids (n = 22 for each alder leaf/metal condition and n = 14 for each chironomid/metal condition) were starved for 2 days and then individually placed in 250 mL beakers filled with Evian^®^ water (Fig. 1). For each metal, they were then either exposed to either two radiolabelled leaf discs for 3-5 h, or to one radiolabelled thawed chironomid larva for 1 h (Fig. 1). For the leaf-feeding condition, the 14 gammarids that ate the most were whole-body gamma-counted alive immediately after the pulse-chase-feeding period. For the larval feeding conditions, all the 14 exposed gammarids were gamma-counted. Selected gammarids, as described below in section 3.1, (i.e. 13 ^109^Cd-leaf-fed gammarids, 11^109^Cd-chironomid-fed gammarids, 13 ^65^Zn-leaf-fed gammarids, 13 ^65^Zn-chironomid-fed gammarids and 14 ^110m^Ag-leaf-fed gammarids) were then transferred under depuration conditions, *i.e.* 200 mL beakers filled with non-radiolabelled with Evian^®^ water (*i.e.* closed circuit, constantly aerated, T = 12.0 ± 0.5°C). Each beaker contained 7 radiolabelled gammarids individually separated by handmade baskets (i.e., plastic mesh 11 cm high and 8.6 cm in diameter, with a mesh size of 0.5 cm; Fig. 1). A non-radiolabelled individual was added to each beaker to control for potential recycling of radioisotopes through to leaching from gammarid depuration. Gammarids were fed *ad libitum* with uncontaminated alder leaves throughout the depuration period (Fig. 1).

**Figure 1.**
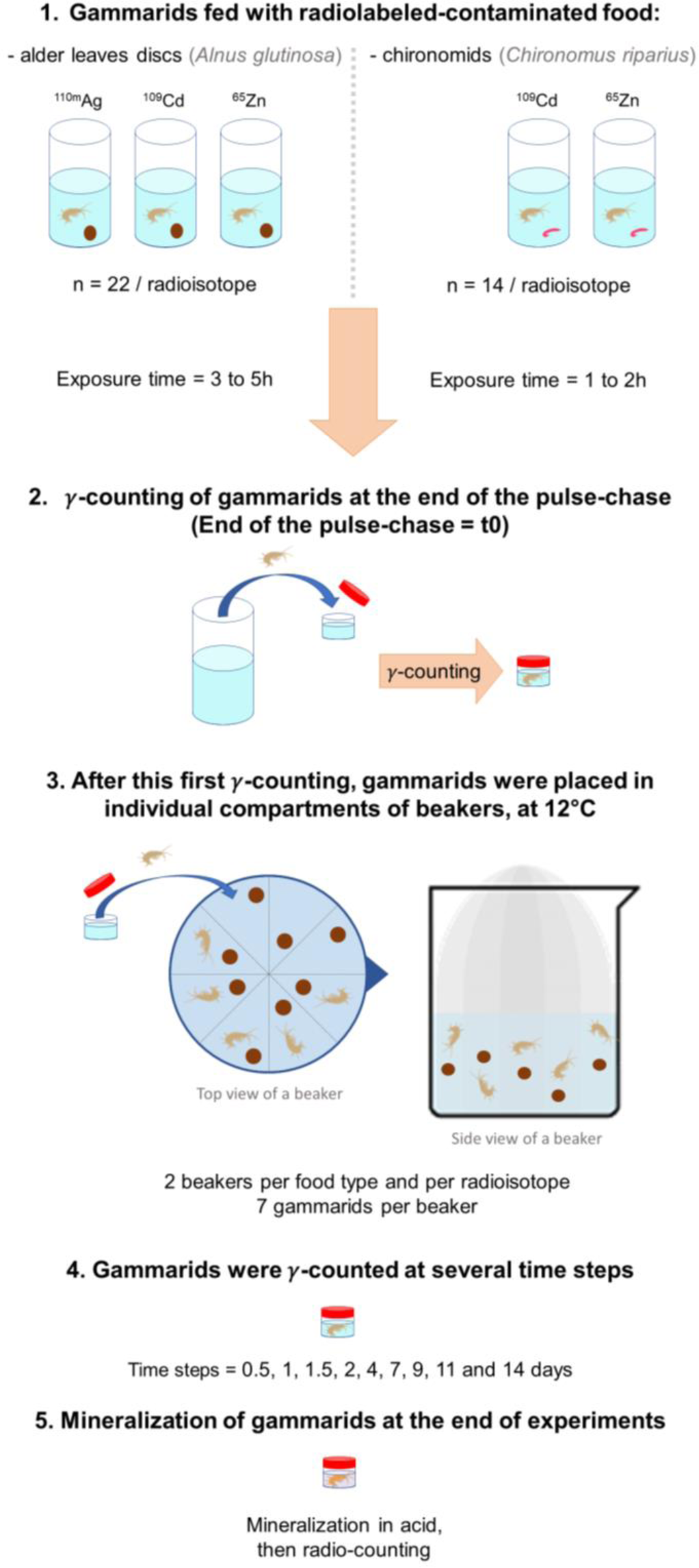
The different steps of the pulse-chase-feeding.

Whole-body radiotracer activities in gammarids were tracked individually during two weeks of depuration (Tables S2 and S3). Individuals were sampled and counted at days 0.5, 1, 1.5, 2, 4, 7, 9, 11 and 14 (Fig. 1). Gammarids were removed from the beakers using a 10 mL pipette, rinsed with clean water and placed in a plastic box (Caubère^®^, ref 1210) containing 500 µL of clean Evian^®^ water (Fig. 1). Counting times of live gammarids varied between 10 and 20 min in order to minimise stress. The counting uncertainties did not exceed 5% for Zn for all samples, whereas the uncertainties increased with decreasing activities in the gammarid samples, without exceeding 20% and 15% for Cd and Ag, respectively. The radiotracer activities in the water were monitored daily and the water was changed at least every two days to minimize tracer contamination in the water. Mortality was also monitored daily.

### 2.5 Estimation of assimilation efficiencies and depuration parameters of gammarids

The depuration of radiotracers by gammarids was expressed as the percentage of remaining activity (i.e. radioactivity at time t divided by the initial radioactivity measured in the organism at the beginning of the depuration period; Tables S2 and S3; see Warnau et al., 1996). The kinetic parameters and the assimilation efficiencies with respect to the food type and metal were estimated using a nonlinear least squares (NLS) approach modelling (see the supplementary material file for the script), following the two-component exponential equation:

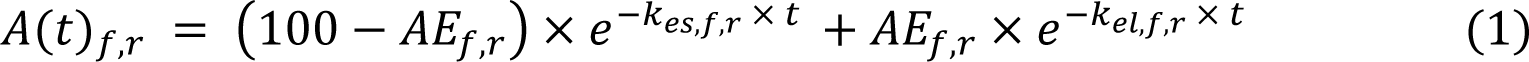

where 𝐴(𝑡)_𝑓,𝑟_is the remaining activity (%) at time 𝑡, with respect to the type of food 𝑓 and the radioisotope 𝑟, 𝐴𝐸_𝑓,𝑟_ is the assimilation efficiency, 𝑘_𝑒𝑠,𝑓,𝑟_ and 𝑘_𝑒𝑙,𝑓,𝑟_ the elimination rates of the first short-term phase (i.e. elimination of the non-assimilated metal fraction, which is rapidly depurated) and of the second long-term phase (i.e. elimination of the assimilated metal fraction, which is slowly depurated), respectively. The indexes 𝑓 correspond to the type of food ingested by the gammarids, with 𝑓 =1 for alder leaves and 𝑓 =2 for chironomid larvae; and the indexes 𝑟 correspond to the radioisotope tested, with 𝑟 =1 for ^109^Cd, 𝑟 =2 for ^65^Zn and 𝑟 =3 for ^110m^Ag. The biological half-life 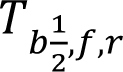 for each metal with respect to the food source was calculated from the elimination rate constant (k_es_ or k_el_), based on the equation:

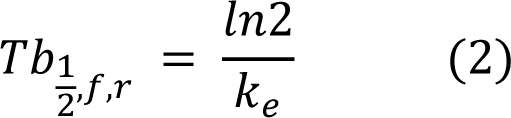

The elimination kinetic parameters and the assimilation efficiencies were estimated using R software.

## 3. Results and discussion

Estimation of metal accumulation efficiencies is an essential first step in understanding bioaccumulation mechanisms. Furthermore, AE values are key inputs for the implementation of toxicokinetic models to study these mechanisms (Van Campenhout et al., 2009). For the gammarid *G. fossarum*, this study provides estimates of the assimilation efficiency and of the two elimination rates of biphasic digestion after the ingestion of food (i.e. of animal or plant origin) contaminated with one of the three metals studied here (Ag, Cd and Zn).

### 3.1 Data quality evaluation and selection

The ^109^Cd is a low energy photon emitter with a gamma ray peak at 88 keV (3.79%), which means that a sample must have enough activity to stand out from the background compared to ^65^Zn (1115 keV) or ^110m^Ag (658 keV for its main gamma ray). Therefore, in order to obtain organisms that have eaten enough and to be able to determine the curve correctly, only individuals that reached a whole-body activity of 150 Bq and above immediately after the single feeding (t0) were kept to follow the metal loss kinetics (Fig. 1 and S1). This means that individuals that did not graze enough during the exposure period due to repletion or stress were dismissed. For the gammarids fed with Cd-enriched chironomids (n=11), only 4 gammarids in a first set of 14 individuals showed a body activity above 150 Bq at t0 (Fig. S1). In a second batch of 14 gammarids, 7 individuals ingested more than 150 Bq of ^109^Cd activities (Fig. S1, Tables S2 and S3). For the Cd leaf-feeding condition (n=13), only 1 of 14 gammarids ingested less than 150 Bq of ^109^Cd at t0, and so was therefore removed from the experiment.

In each condition (n=13) of dietary Zn exposure (i.e. both leaves and larvae as sources), one gammarid died during the first day of the experiment. In the leaf-feeding Ag condition (n=14), all 14 gammarids remained alive throughout the depuration phase.

### 3.2. Metal elimination kinetics patterns

Kinetic models were fitted to whole body concentrations over time following a pulse-chase-feeding of metal-contaminated alder leaves. The metal elimination kinetics showed a biphasic pattern (Fig. 2), expressing the presence of two metal pools: a first poorly retained pool characterised by a high rate of elimination (k_es_), and a second slowly eliminated pool (k_el_).

**Figure 2.**
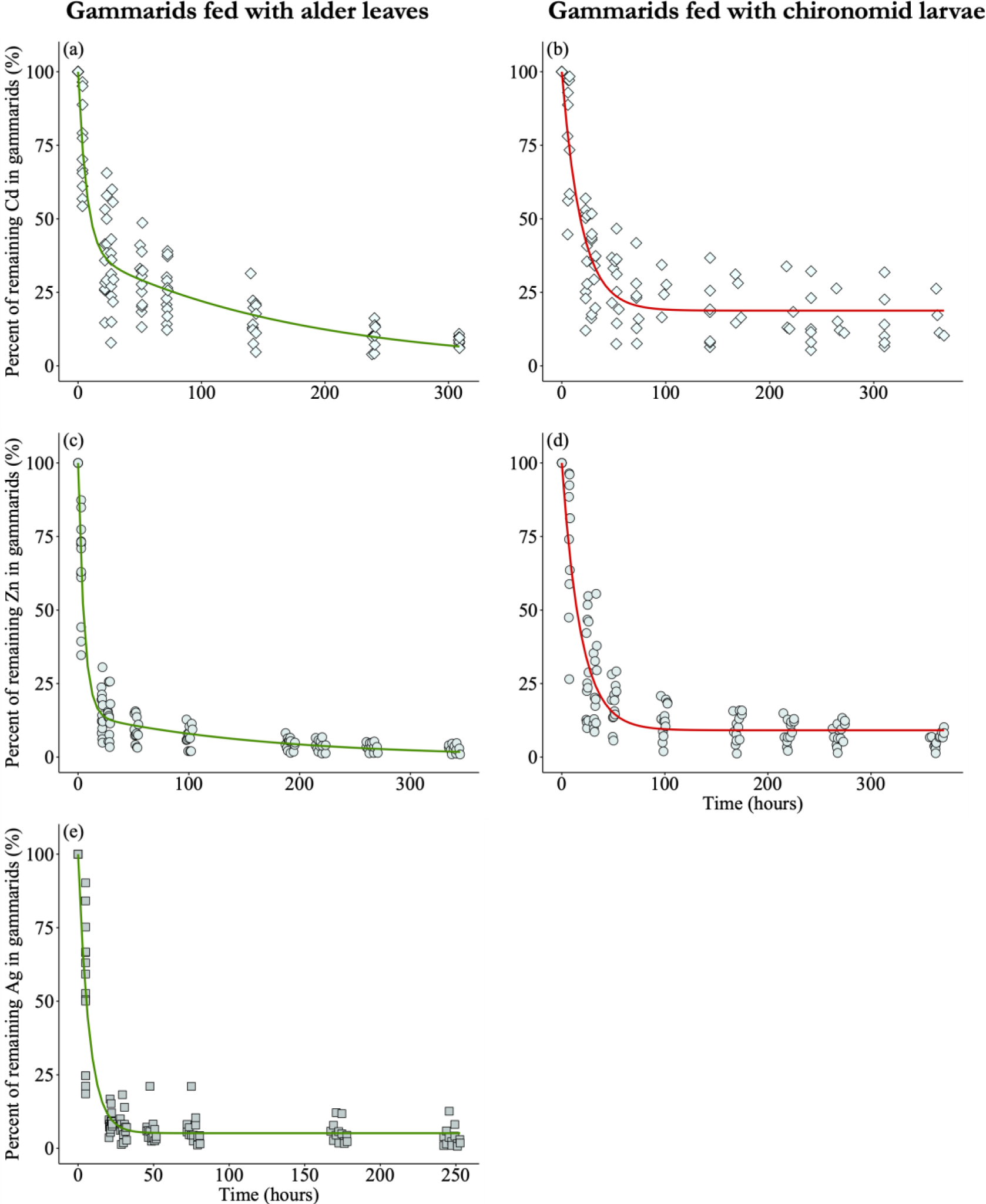
Influence of the metal and the type of matrix used for feeding on the elimination of metals ingested by the gammarids during the depuration phase, expressed as the percentage of activity remaining in the gammarids compared to the end of the pulse-chase phase (t0) as a function of the depuration time (hours). Gammarids were contaminated by cadmium for a) and b); zinc for c) and d); and silver for e). Food corresponded to leaf discs for a), c) and e); and to chironomid larvae for b) and d).

Regardless of the metal, gammarids eliminated the highest proportions of ingested trace elements during the first 48 h after exposure to contaminated leaves. Elimination rates were similar among metals with k_es_ values of 0.18 ± 0.01 h^-1^, 0.15 ± 0.03 h^-1^ and 0.14 ± 0.01 h^-1^ for Zn, Cd and Ag respectively (Table 1). Thus, these values imply that, after a 5 h period of grazing, these metal pools had a Tb_s1/2_ of 4-5 hours (Table 1). These biological half-lives, which are similar among metals, are consistent with the intestinal transit time reported for *Gammarus pulex* (16 hours; Pellet et al., 2014), suggesting that these fractions were not assimilated and were excreted in feces. This hypothesis, stimulated by the ingestion of non-radiolabelled alder leaves since the gammarids were placed in depuration conditions, is supported by this significant release of radiotracers during the first 48 hours of the elimination phase.

**Table 1.** Parameters estimated by fitting an NLS model from pulse-chase feeding data, with EA (%), kel (d^-1^) and kes (d^-1^) with their respective standard deviations, t-value, Pr(>ItI) and percentiles (95%). Calculated biological half-lives times of metals as a function of diet type with their respective standard deviations and percentiles (95%).

| Metal | Food type | Parameters | Assimilation efficiencies |  |  |  |  | Biological half-life |  |  |
| --- | --- | --- | --- | --- | --- | --- | --- | --- | --- | --- |
| | | | Estimate values | t value | $\text{Pr}(>\vert t\vert )$ | Percentiles | | Calculated values | Percentiles | |
| Ag | Alder leaves | $k_{es}$ | $0.14 \pm 0.01$ | 14 | <2E-16 | 0.12 | 0.16 | $5.1 \pm 0.4$ | 4.4 | 5.9 |
| | | AE | $5.2 \pm 1.1$ | 4.8 | 4.2E-6 | 3.0 | 7.3 | / | / | / |
| | | $k_{el}$ | 0 | / | / | / | / | ∞ | / | / |
| Cd | Chironomid larvae | $k_{es}$ | $0.053 \pm 0.005$ | 11 | <2E-16 | 0.044 | 0.063 | $13 \pm 1$ | 11 | 16 |
| | | AE | $19 \pm 2$ | 11 | <2E-16 | 15 | 22 | / | / | / |
| | | $k_{el}$ | 0 | / | / | / | / | ∞ | / | / |
| | Alder leaves | $k_{es}$ | $0.15 \pm 0.03$ | 5.5 | 3.0e-7 | 0.09 | 0.20 | $4.9 \pm 1.0$ | 3.6 | 7.2 |
| | | AE | $39 \pm 3$ | 12 | <2E-16 | 33 | 45 | / | / | / |
| | | $k_{el}$ | $0.0058 \pm 0.0010$ | 5.6 | 1.8e-7 | 0.0037 | 0.0078 | $124 \pm 25$ | 89 | 185 |
| Zn | Chironomid larvae | $k_{es}$ | $0.055 \pm 0.004$ | 15 | <2E-16 | 0.047 | 0.062 | $13 \pm 1$ | 11 | 15 |
| | | AE | $9.1 \pm 1.2$ | 7.3 | 2.6E-11 | 6.7 | 12 | / | / | / |
| | | $k_{el}$ | 0 | / | / | / | / | ∞ | / | / |
| | Alder leaves | $k_{es}$ | $0.18 \pm 0.01$ | 13 | <2E-16 | 0.16 | 0.21 | $3.8 \pm 0.3$ | 3.3 | 4.4 |
| | | AE | $15 \pm 2$ | 8.4 | 8.0E-14 | 11 | 18 | / | / | / |
| | | $k_{el}$ | $0.0061 \pm 0.0014$ | 4.4 | 2.3E-5 | 0.0034 | 0.0089 | $120 \pm 35$ | 78 | 203 |

The second kinetic phase was characterised by a lower elimination rate, with k_el_ values of 0.0061 ± 0.0014 h^-1^, 0.0058 ± 0.0010 h^-1^ and zero for Zn, Cd and Ag, corresponding to k_el_ of Tb_l1/2_ = 120 ± 35 h, 124 ± 25 h and ∞, respectively (Table 1). This longer elimination rate is considered to be due to the physiological turnover of the assimilated metal fraction. It is assumed that this pool crosses the intestinal barrier, is distributed to different organs and is taken over by detoxification mechanisms, as already described for Cd and Zn in invertebrates (Nunez-Nogueira et al., 2006). It is noteworthy that the Tb_l1/2_ of Cd and Zn are equivalent, suggesting that metabolism of both trace elements requires similar physiological mechanisms. In contrast, the assimilated pool of Ag was not eliminated by the gammarid, which contrasts with a shorter Ag retention time than other metals, such as Cd or Zn, reported in daphnia (Lam & Wang, 2006), in cuttlefish and in turbot (Bustamante et al., 2004; Pouil et al., 2015). This phenomenon has already been observed in *Lymnea stagnalis* (k_el_ not different from 0) suggesting that Ag is strongly retained in internal tissues (Croteau et al., 2011). This could be explained by the fact that Ag is an element that reacts very strongly with sulphur compounds (Lam & Wang, 2006). In bivalves, Ag has indeed been shown to be stored in the epithelial cells of the intestinal cells, as Ag_2_S (Berthet et al., 1992), which are already known to be a site of accumulation and detoxification of other metals such as Cd and Cu in gammarids (Pellet et al., 2014). In this highly stable Ag_2_S form, silver is found in the insoluble fraction (Berthet et al., 1992), and remains indefinitely sequestered in tissues in a non-metabolically available form (Fowler et al., 2004).

### 3.3. The AE values of metals after feeding by contaminated alder leaves

According to the second depuration kinetic components, the assimilation efficiencies of metals in alder leaves fed to gammarids were ranked as follows: Cd (39 ± 3%) > Zn (15 ± 2%) > Ag (5.2 ± 1.1%) (Fig. 2; Table 1). These contrasting AE values among elements were also found in *Daphnia magna*, with AE of 30-70%, 7-66% and 1-2%, for Cd, Zn and Ag, respectively (Yu & Wang, 2002; Lam & Wang, 2006). Consistent with our results, a higher AE for Cd (5-47%) than for Zn (1.4%) was also found in the leaf-fed gammarid *Gammarus pulex* (Xu & Pascoe, 1994; Pellet et al., 2014). To our knowledge, there are no data on Ag AE in amphipods, although similar AE values ranging from 3 to 23% have been reported in the marine copepods *Acartia torua*, *Acartia hudsonica* and *Temova longicornis*. Compared to the Ag AE determined here in *G. fossarum*, higher AE values were reported for this metal in the cephalopod *Sepia officinalis* (19 ± 3%) (Bustamante et al., 2004) and in the gastropod *Lymnaea stagnalis* (73 ± 5%) (Croteau et al., 2011), whereas it did not exceed 1% in the fish *Scophthalmus maximus* (0.32%) (Reinfelder & Fisher, 1991; Lam & Wang, 2006; Pouil et al., 2015). Thus, all these data show large disparities confirming that there are no general rules for predicting AE among metals and biological species (Wang & Fisher, 1999a; Pouil et al., 2018).

### 3.4. The effect of food type on metal trophic transfer

The AE values are intrinsically linked to both biotic and abiotic factors that modulate the digestive physiology, metal bioavailability and bioaccessibility (Pouil et al., 2018). Among these factors, the type of food and its amount significantly influence the AE values, as reported in fish (Pouil et al., 2017), while it is less evident in copepods, as reviewed by Wang & Fisher (1999b). More specifically, the AE of Cd in *Dreissena polymorpha* has been estimated to range from 19 to 72% for 8 types of food (Roditi & Fisher, 1999). The AE of Zn has also been reported to be highly variable in fish, ranging from 2 to 97% (with a median of 20%) as a function of the predator species and the food sources (Pouil et al., 2018). Mechanistically, these AE variations are thought to be driven by the subcellular distribution of metals in food (i.e. the storage form). Theoretically, metals are considered to be trophically available for transfer to consumers when they are associated with organelles, enzymes and metal-binding proteins (i.e. Trophically Available Metals, abbreviated TAM), i.e. metals found in the soluble fraction. Conversely, metals associated with the insoluble fraction (i.e. metal-rich granules or cellular debris) may not be assimilated (Wallace & Lopez, 1996; Wallace & Luoma, 2003). Nevertheless, in practice, the relationship between TAM and AE is less clear due to the complex organ and subcellular distribution of metals in metazoan prey (see Pouil et al., 2016).

In the present study, the AE of Cd and Zn were 2- and 1.5-fold higher when the ingested food was alder leaves (39 ± 3% and 15 ± 2%, respectively; Fig. 2 and Table 1) compared to chironomid larvae (19 ± 2% and 9.1 ± 1.2%, respectively; Fig. 2 and Table 1). This is in contrast to what is usually reported for omnivorous organisms, where the trophic transfer of metals from plant sources is lower than that from animal prey because the digestion process of plant materials is slower and more complex (Xu & Pascoe, 1994; Nunez-Nogueira et al., 2006). For example, the shrimp *Penaeus indicus* fed with Cd-contaminated algae showed an AE of 42.4 ± 5.1%, a value 1.8 time lower than that reported for shrimp fed with Cd-contaminated squid (*i.e.* 74.6 ± 8.5%). In contrast, the AE of Zn was not dependent on the type of diet (i.e. AE of *P. indicus* fed with squid = 57.9 ± 7.3%; AE of *P. indicus* fed with algae = 59.1 ± 8.4%) (Nunez-Nogueira et al., 2006).

Regarding our results, it is noteworthy that live chironomid larvae were contaminated with the radiotracers whereas only dead leaves were radiolabelled. More than the plant or animal character, we suspect a large passive adsorption of metals on the walls of inert dead leaves compared to a poor absorption within the leaf structures. Once ingested, the metal fraction should be easily desorbed under a free form ion and thus more bioavailable, increasing the assimilation efficiency for the gammarid. On the contrary, metals actively ingested by live chironomid larvae were distributed among tissues, incorporated into cells and taken up by detoxification mechanisms, thus binding to various subcellular components. Therefore, the higher AE obtained for leaf-fed gammarids than for chironomid-fed gammarids could be related to the sorption processes and the TAM fractions in live vs. inert food sources rather than to the contrasting efficiencies of digestion of plant and animal materials.

As found for AE, both k_es_ and k_el_ of Cd and Zn varied more with the food matrix than with the metal itself. With Tb_s1/2_ of 13.0 ± 1.2 and 13.0 ± 0.9 h, for Cd and Zn, respectively, the short-lived metal pool was 2 to 3 times longer in chironomid-fed gammarids than in leaf-fed gammarids (Table 1). In addition, the long-term biological half-life Tb_l1/2_ tended to infinity for both elements when gammarids were fed with chironomids (Table 1). This strong retention of metals is consistent with the previously stated hypothesis that the two food sources have different TAM (Rainbow et al., 2011) which, once assimilated, should also determine the fate of the metal in the gammarid. Thus, the leaf-adsorbed metal assimilated in a free form or in a weak ionic complex is likely to be taken up more rapidly by detoxification and/or elimination processes than the assimilated metal strongly bound to components and concretions of chironomid cells. Nunez-Nogeira et al (2006) also found a higher k_e_ of Cd in the shrimp *P. indicus* contaminated with squid meat (0.009 ± 0.003 d^-1^) than when contaminated with algae (0.004 ± 0.001 d^-1^), confirming that Cd assimilated from a vegetable matrix is eliminated more rapidly than from a complex multicellular organism. However, Zn was excreted more rapidly after ingestion of animal than plant matrices (Nunez-Nogueira et al., 2006), contrary to what we found in *G. fossarum*. These observations suggest that Zn and Cd undergo different physiological bioaccumulation processes depending on the type of food ingested, hence the importance of better understanding the toxicokinetics of ingested metals and what influences it.

## 5. Conclusion

To better improve and predict the fate of metals in organisms and their potential transfer within food webs, it is important to understand the key factor controlling metal bioaccumulation in organisms, such as diet as a contamination pathway (Xu & Pascoe, 1994; Hadji et al., 2016). As a first step, the results presented here fill in the gaps regarding the critical physiological parameters governing bioaccumulation and trophic transfer mechanisms: assimilation efficiency (AE) and elimination rates. Furthermore, AE is a key input for the implementation of toxicokinetic models to study bioaccumulation mechanisms and the fate of metals in organisms (Van Campenhout et al., 2009). By exposing the gammarid, *G. fossarum*, to two types of food (i.e. of animal or plant origin) contaminated with Ag, Cd and Zn, this study reported that, regardless of the type of food, Cd had the highest AE values (39 and 19%, respectively), followed by Zn (15 and 9%, respectively) and Ag (5%, determined for plant only). In turn, the food matrix modulated the metal elimination rates, as k_es_ and k_el_ for Cd and Zn were higher when metals were ingested *via* leaves than *via* chironomids. This implies a shorter retention time for metals presumably adsorbed on an inert matrix compared to metals metabolically assimilated by live chironomids.

Thus, by providing AE estimates, we have shown that AE in gammarids depends on metals and food sources, as has already been demonstrated in fish (Pouil et al., 2018) or marine invertebrates (Bustamante et al., 2004). It is well known that the metal subcellular partitioning and its storage form in the food source is an important parameter affecting the AE of predators. Therefore, it would be interesting to demonstrate the relationship between bioavailable metal fractions in food (e.g. TAM, Rainbow et al., 2011) and AE in order to better predict trophic transfer based on food type.

Finally, the use of AE data could greatly improve toxicokinetic models to better understand the ADME (i.e. Absorption, Distribution, Metabolisation and Elimination) mechanisms of metals in small invertebrates. In particular, the data obtained here has enabled us to improve our knowledge and set up a toxicokinetic model of Cd accumulation *via* the trophic pathway (Gestin et al., *submitted*). The study of organotropism over time in relation to the metal accumulation pathway in gammarids would be an important direction for future research to further unravel the bioaccumulation mechanisms (Franklin et al., 2005; Urien et al., 2016) and to implement toxicokinetic models.

## Acknowledgments

This work has been supported by the APPROve project funded by the ANR (ANR-18-CE34-0013-01). We thank the « Radioecology lab » of the LIENSs and Competent Radiological Protection Persons (Christine Dupuy and Thomas Lacoue-Labarthe) for their technical support. This work benefitted from the French GDR “Aquatic Ecotoxicology” framework which aims at fostering stimulating scientific discussions and collaborations for more integrative approaches.

