## Supplemental files for "Assimilation efficiencies and elimination rates of silver, cadmium and zinc accumulated by trophic pathway in *Gammarus fossarum*"

Table S1. Characteristics of waters used for the experiments: well water to leach the leaves and Évian® water in all other cases.

| | Well water | Évian® |
|---|---|---|
| Bicarbonates $HCO_3^-$ (mg.L$^{-1}$) | 209 | 360 |
| Calcium $Ca^{2+}$ (mg.L$^{-1}$) | 73.7 | 80 |
| Chlorides $Cl^-$ (mg.L$^{-1}$) | 11.5 | 10 |
| Magnesium $Mg^{2+}$ (mg.L$^{-1}$) | 7.3 | 26 |
| Nitrates $NO_3^-$ (mg.L$^{-1}$) | 6.5 | 3.8 |
| Potassium $K^+$ (mg.L$^{-1}$) | 1.9 | 1 |
| Silica $SiO_2$ (mg.L$^{-1}$) | 6.1 | 15 |
| Sodium $Na^+$ (mg.L$^{-1}$) | 8.2 | 6.5 |
| Sulfates $SO_4^{2-}$ (mg.L$^{-1}$) | 35.9 | 14 |
| pH | 7.55 | 7.2 |

Table S2. Measured amount of $^{109}$Cd (Bq) in gammarids along the depuration phase in function of the metal and the type of food tested, with t0 is the end of the feeding phase.

| Radioisotope<br>Type of food | Gammarid | Time (hours) | | | | | | | |
|---|---|---|---|---|---|---|---|---|---|
| | | 0 | 5.0 | 21 | 28 | 46 | 72 | 167 | 242 |
| Ag<br>Alder leaves | 1 | 171 | 42 | 17 | 11 | 12 | 14 | 10 | 6.7 |
| | 2 | 155 | 117 | 5.6 | 16 | 9.2 | 8.6 | | 1.5 |
| | 3 | 208 | 131 | 17 | 17 | 13 | 14 | 16 | 12 |
| | 4 | 192 | 101 | 14 | 2.5 | 7.1 | 8.9 | 5.1 | 2.9 |
| | 5 | 284 | 143 | 21 | 18 | 16 | 13 | 13 | 11 |
| | 6 | 129 | 117 | 22 | 24 | 27 | 27 | 16 | 16 |
| | 7 | 219 | 46 | 12 | 4.1 | 5.3 | 5.4 | 3.9 | 2.2 |
| | 8 | 233 | 58 | 15 | 11 | 8.2 | 10 | 13 | 5.3 |
| | 9 | 156 | 78 | 24 | 22 | 13 | 16 | 18 | 13 |
| | 10 | 129 | 86 | 12 | 11 | 4.8 | 10 | 6.4 | 5.4 |
| | 11 | 185 | 123 | 17 | 13 | 12 | 7.4 | 3.1 | 1.3 |
| | 12 | 198 | 117 | 24 | 11 | 5.0 | 2.2 | 4.5 | 3.1 |
| | 13 | 219 | 184 | 16 | 6.0 | 7.0 | 3.3 | 5.0 | 4.0 |
| | 14 | 284 | 53 | 24 | 20 | 12 | 12 | 13 | 8.4 |

| Radioisotope<br>Type of food | Gammarid | 0 | 3.6 | 7.5 | 23 | 24 | 27 | 31 | 48 | 52 | 72 | 96 | 143 | 167 | 216 | 240 | 264 | 309 | 360 |
|---|---|---|---|---|---|---|---|---|---|---|---|---|---|---|---|---|---|---|---|
| Cd<br>Chironomid larvae | 1 | 170 | | 99 | | 69 | | 62 | 62 | | | 58 | | 53 | 57 | | 45 | | 45 |
| | 2 | 180 | | 177 | | 64 | | 53 | 39 | | | 30 | | 26 | 24 | | 27 | | 31 |
| | 3 | 150 | | 110 | | 42 | | 51 | 50 | | | 36 | | 42 | 19 | | 18 | | 17 |
| | 4 | 165 | | 160 | | 84 | | 62 | 59 | | | 46 | | 27 | 30 | | 18 | | 17 |
| | 5 | 396 | 223 | | 100 | | 78 | | | 76 | 63 | | 25 | | | 21 | | | |
| | 6 | 309 | 287 | | 164 | | 139 | | | 96 | 71 | | 59 | | | 39 | | 31 | |
| | 7 | 272 | 265 | | 137 | | 117 | | | 127 | 114 | | 100 | | | 88 | | 87 | |
| | 8 | 275 | 244 | | 63 | | 48 | | | 40 | 35 | | 23 | | | 22 | | 22 | |
| | 9 | 164 | 73 | | 20 | | 27 | | | 12 | 12 | | 13 | | | | | 23 | |
| | 10 | 164 | 128 | | 93 | | 85 | | | 59 | 46 | | 42 | | | 38 | | 37 | |
| | 11 | 171 | 177 | | 89 | | 75 | | | 43 | 41 | | 31 | | | 20 | | 11 | |

| Radioisotope<br>Type of food | Gammarid | 0 | 3.6 | 23 | 27 | 52 | 72 | 143 | 240 | 309 |
|---|---|---|---|---|---|---|---|---|---|---|
| Cd<br>Alder leaves | 1 | 330 | 232 | 118 | 103 | 83 | 69 | 43 | 33 | 29 |
| | 2 | 257 | 146 | 66 | 70 | 71 | 59 | 33 | 10 | |
| | 3 | 234 | 238 | 97 | 84 | 71 | 71 | 33 | | 23 |
| | 4 | 214 | 116 | 56 | 63 | 45 | 30 | 26 | | |
| | 5 | 765 | 509 | 407 | 60 | 248 | 221 | 151 | 80 | 83 |
| | 6 | 285 | 225 | 81 | 68 | 52 | 35 | 21 | 12 | 22 |
| | 7 | 168 | 168 | 25 | 37 | 22 | 32 | 8 | | 11 |
| | 8 | 573 | 375 | 167 | 86 | 116 | 97 | 64 | 41 | 35 |
| | 9 | 196 | 189 | 98 | | 65 | 52 | 35 | 20 | 19 |
| | 10 | 203 | 180 | 133 | 113 | 99 | 76 | 45 | 27 | |
| | 11 | 344 | 327 | 199 | 206 | 141 | 134 | 108 | | |
| | 12 | 206 | 126 | 85 | 89 | 80 | 53 | 42 | 28 | 17 |
| | 13 | 182 | 141 | 70 | 70 | 59 | 69 | 38 | 30 | 18 |

| Radioisotope<br>Type of food | Gammarid | 0 | 6.8 | 24 | 31 | 49 | 97 | 167 | 216 | 264 | 359 |
|---|---|---|---|---|---|---|---|---|---|---|---|
| Zn<br>Chironomid larvae | 1 | 170 | 81 | 22 | 19 | 33 | 22 | 15 | 11 | 12 | 12 |
| | 2 | 122 | 162 | 52 | 43 | 34 | 25 | 19 | 18 | 12 | 8.2 |
| | 3 | 785 | 581 | 94 | 76 | 54 | 39 | 33 | 27 | 25 | 22 |
| | 4 | 463 | 409 | 104 | 60 | 62 | 40 | 36 | 31 | 30 | 22 |
| | 5 | 437 | 116 | 43 | 37 | 24 | 8.5 | 5.2 | 9.3 | 6.2 | 5.8 |
| | 6 | 576 | 556 | 145 | 117 | 79 | 41 | 32 | 31 | 24 | 22 |
| | 7 | 219 | 202 | 102 | 71 | 46 | 31 | 22 | 15 | 14 | 15 |
| | 8 | 317 | 186 | 164 | 55 | 49 | 38 | 41 | 34 | 30 | 22 |
| | 9 | 188 | 181 | 44 | 37 | 27 | 20 | 11 | 16 | 10 | 13 |
| | 10 | 193 | 156 | 105 | 107 | 47 | 38 | 31 | 23 | 26 | 17 |
| | 11 | 71 | 45 | 8.8 | 8.3 | 14 | 8.6 | 8.1 | 9.1 | 8.0 | 2.8 |
| | 12 | 175 | 142 | 81 | 66 | 51 | 32 | 25 | 21 | 19 | 18 |
| | 13 | 140 | 167 | 40 | 41 | 31 | 25 | 22 | 18 | 17 | 11 |

| Radioisotope<br>Type of food | Gammarid | 0 | 2.8 | 21 | 27 | 51 | 98 | 187 | 215 | 260 | 335 |
|---|---|---|---|---|---|---|---|---|---|---|---|
| Zn<br>Alder leaves | 1 | 1036 | 802 | 202 | 161 | 97 | 60 | 44 | 41 | 35 | 34 |
| | 2 | 1395 | 1184 | 331 | 209 | 207 | 178 | 114 | 91 | 72 | 56 |
| | 3 | 1220 | 768 | 149 | 165 | 90 | 72 | 37 | 35 | 34 | 29 |
| | 4 | 2134 | 943 | 174 | 113 | 88 | 43 | 45 | 39 | 28 | 20 |
| | 5 | 943 | 668 | 201 | 242 | 147 | 75 | 60 | 54 | 45 | 43 |
| | 6 | 1288 | 942 | 183 | 174 | 97 | 81 | 71 | 48 | 46 | 36 |
| | 7 | 915 | 559 | 182 | 129 | 139 | 74 | 52 | 49 | 45 | 34 |
| | 8 | 873 | 343 | 53 | 45 | 29 | 17 | 13 | 10 | 12 | 10 |
| | 9 | 802 | 501 | 79 | 61 | 64 | 52 | 44 | 31 | 27 | 20 |
| | 10 | 593 | 435 | 181 | 152 | 81 | 62 | 39 | 40 | 31 | 28 |
| | 11 | 919 | 319 | 45 | 31 | 29 | 19 | 17 | 12 | 13 | 8.3 |
| | 12 | 997 | 871 | 176 | 180 | 110 | 94 | 49 | | | |
| | 13 | 825 | 597 | 100 | 94 | 62 | 94 | 34 | 33 | 29 | 24 |

Table S3. Percent of remaining metal in gammarids compared to t0 (%) along the depuration phase in function of the metal and the type of food tested, with t0 is the end of the feeding phase.

**Ag – Alder leaves** (Time, hours)

| Radioisotope / Type of food | Gammarid | 0 | 5.0 | 21 | 28 | 46 | 72 | 167 | 242 |
|---|---|---|---|---|---|---|---|---|---|
| Ag Alder leaves | 1 | 100 | 25 | 10 | 6.1 | 7.2 | 8.1 | 5.8 | 3.9 |
| | 2 | 100 | 75 | 3.6 | 10 | 5.9 | 5.6 | | 0.9 |
| | 3 | 100 | 63 | 8.2 | 8.2 | 6.1 | 7.0 | 7.8 | 5.9 |
| | 4 | 100 | 53 | 7.2 | 1.3 | 3.7 | 4.6 | 2.7 | 1.5 |
| | 5 | 100 | 50 | 7.4 | 6.5 | 5.7 | 4.5 | 4.5 | 3.7 |
| | 6 | 100 | 90 | 17 | 18 | 21 | 21 | 12 | 13 |
| | 7 | 100 | 21 | 5.4 | 1.9 | 2.4 | 2.5 | 1.8 | 1.0 |
| | 8 | 100 | 25 | 6.5 | 4.6 | 3.5 | 4.5 | 5.7 | 2.3 |
| | 9 | 100 | 50 | 15 | 14 | 8 | 10 | 12 | 8.1 |
| | 10 | 100 | 67 | 9.2 | 8.2 | 3.7 | 7.9 | 4.9 | 4.2 |
| | 11 | 100 | 67 | 9.1 | 7.2 | 6.4 | 4.0 | 1.7 | 0.7 |
| | 12 | 100 | 59 | 12 | 5.8 | 2.5 | 1.1 | 2.3 | 1.6 |
| | 13 | 100 | 84 | 7.5 | 2.7 | 3.2 | 1.5 | 2.3 | 1.8 |
| | 14 | 100 | 19 | 8.3 | 7.1 | 4.1 | 4.3 | 4.4 | 3.0 |

**Cd – Chironomid larvae** (Time, hours)

| Radioisotope / Type of food | Gammarid | 0 | 3.6 | 7.5 | 23 | 24 | 27 | 31 | 48 | 52 | 72 | 96 | 143 | 167 | 216 | 240 | 264 | 309 | 360 |
|---|---|---|---|---|---|---|---|---|---|---|---|---|---|---|---|---|---|---|---|
| Cd Chironomid larvae | 1 | 100 | | 58 | | 41 | | 37 | 37 | | | 34 | | 31 | 34 | | 26 | | 26 |
| | 2 | 100 | | 98 | | 36 | | 29 | 21 | | | 17 | | 15 | 13 | | 15 | | 17 |
| | 3 | 100 | | 73 | | 28 | | 34 | 33 | | | 24 | | 28 | 13 | | 12 | | 11 |
| | 4 | 100 | | 97 | | 51 | | 37 | 36 | | | 28 | | 16 | 18 | | 11 | | 10 |
| | 5 | 100 | 56 | | 25 | | 20 | | | 19 | 16 | | 6.4 | | | 5.4 | | | |
| | 6 | 100 | 93 | | 53 | | 45 | | | 31 | 23 | | 19 | | | 13 | | 10 | |
| | 7 | 100 | 97 | | 50 | | 43 | | | 47 | 42 | | 37 | | | 32 | | 32 | |
| | 8 | 100 | 89 | | 23 | | 17 | | | 14 | 13 | | 8.3 | | | 8.0 | | 7.9 | |
| | 9 | 100 | 45 | | 12 | | 16 | | | 7.5 | 7.6 | | 7.9 | | | | | 14 | |
| | 10 | 100 | 78 | | 57 | | 52 | | | 36 | 28 | | 26 | | | 23 | | 23 | |
| | 11 | 100 | 104 | | 52 | | 44 | | | 25 | 24 | | 18 | | | 11 | | 6.6 | |

**Cd – Alder leaves** (Time, hours)

| Radioisotope / Type of food | Gammarid | 0 | 3.6 | 23 | 27 | 52 | 72 | 143 | 240 | 309 |
|---|---|---|---|---|---|---|---|---|---|---|
| Cd Alder leaves | 1 | 100 | 70 | 36 | 31 | 25 | 21 | 13 | 10 | 8.8 |
| | 2 | 100 | 57 | 26 | 27 | 28 | 23 | 13 | 3.9 | |
| | 3 | 100 | 102 | 41 | 36 | 31 | 30 | 14 | | 10 |
| | 4 | 100 | 54 | 26 | 29 | 21 | 14 | 12 | | |
| | 5 | 100 | 67 | 53 | 8 | 32 | 29 | 20 | 10 | 11 |
| | 6 | 100 | 79 | 28 | 24 | 18 | 12 | 7.5 | 4.2 | 7.7 |
| | 7 | 100 | 100 | 15 | 22 | 13 | 19 | 4.7 | | 6.3 |
| | 8 | 100 | 65 | 29 | 15 | 20 | 17 | 11 | 7.2 | 6.0 |
| | 9 | 100 | 96 | 50 | | 33 | 27 | 18 | 10 | 10 |
| | 10 | 100 | 89 | 66 | 56 | 49 | 37 | 22 | 13 | |
| | 11 | 100 | 95 | 58 | 60 | 41 | 39 | 31 | | |
| | 12 | 100 | 61 | 41 | 43 | 39 | 26 | 21 | 14 | 8.2 |
| | 13 | 100 | 77 | 38 | 38 | 32 | 38 | 21 | 16 | 10 |

**Zn – Chironomid larvae** (Time, hours)

| Radioisotope / Type of food | Gammarid | 0 | 6.8 | 24 | 31 | 49 | 97 | 167 | 216 | 264 | 359 |
|---|---|---|---|---|---|---|---|---|---|---|---|
| Zn Chironomid larvae | 1 | 100 | 47 | 13 | 11 | 19 | 13 | 8.5 | 6.8 | 6.8 | 6.9 |
| | 2 | 100 | 132 | 42 | 35 | 28 | 21 | 16 | 15 | 10 | 6.7 |
| | 3 | 100 | 74 | 12 | 10 | 6.9 | 4.9 | 4.2 | 3.4 | 3.2 | 2.8 |
| | 4 | 100 | 88 | 22 | 13 | 13 | 8.7 | 7.7 | 6.8 | 6.4 | 4.8 |
| | 5 | 100 | 27 | 10 | 8.6 | 5.6 | 2.0 | 1.2 | 2.1 | 1.4 | 1.3 |
| | 6 | 100 | 96 | 25 | 20 | 14 | 7.1 | 5.6 | 5.3 | 4.2 | 3.7 |
| | 7 | 100 | 92 | 47 | 33 | 21 | 14 | 10 | 6.9 | 6.4 | 6.7 |
| | 8 | 100 | 59 | 52 | 17 | 16 | 12 | 13 | 11 | 9.5 | 7.0 |
| | 9 | 100 | 96 | 24 | 20 | 15 | 10 | 5.9 | 8.4 | 5.2 | 6.7 |
| | 10 | 100 | 81 | 55 | 56 | 24 | 20 | 16 | 12 | 13 | 8.6 |
| | 11 | 100 | 64 | 12 | 12 | 19 | 12 | 11 | 13 | 11 | 3.9 |
| | 12 | 100 | 81 | 46 | 38 | 29 | 19 | 15 | 12 | 11 | 10 |
| | 13 | 100 | 119 | 29 | 30 | 22 | 18 | 16 | 13 | 12 | 8.2 |

**Zn – Alder leaves** (Time, hours)

| Radioisotope / Type of food | Gammarid | 0 | 2.8 | 21 | 27 | 51 | 98 | 187 | 215 | 260 | 335 |
|---|---|---|---|---|---|---|---|---|---|---|---|
| Zn Alder leaves | 1 | 100 | 77 | 20 | 16 | 9.3 | 5.8 | 4.3 | 3.9 | 3.4 | 3.3 |
| | 2 | 100 | 85 | 24 | 15 | 15 | 13 | 8.2 | 6.5 | 5.2 | 4.0 |
| | 3 | 100 | 63 | 12 | 13 | 7.3 | 5.9 | 3.1 | 2.9 | 2.8 | 2.4 |
| | 4 | 100 | 44 | 8.2 | 5.3 | 4.1 | 2.0 | 2.1 | 1.8 | 1.3 | 0.9 |
| | 5 | 100 | 71 | 21 | 26 | 16 | 7.9 | 6.4 | 5.7 | 4.7 | 4.6 |
| | 6 | 100 | 73 | 14 | 14 | 7.5 | 6.3 | 5.5 | 3.7 | 3.6 | 2.8 |
| | 7 | 100 | 61 | 20 | 14 | 15 | 8.1 | 5.7 | 5.3 | 4.9 | 3.7 |
| | 8 | 100 | 39 | 6.1 | 5.1 | 3.3 | 2.0 | 1.5 | 1.2 | 1.4 | 1.1 |
| | 9 | 100 | 62 | 10 | 7.6 | 8.0 | 6.4 | 5.4 | 3.8 | 3.4 | 2.5 |
| | 10 | 100 | 73 | 31 | 26 | 14 | 10 | 6.6 | 6.8 | 5.3 | 4.8 |
| | 11 | 100 | 35 | 4.9 | 3.4 | 3.2 | 2.0 | 1.8 | 1.3 | 1.4 | 0.9 |
| | 12 | 100 | 87 | 18 | 18 | 11 | 9.4 | 4.9 | | | |
| | 13 | 100 | 72 | 12 | 11 | 7.5 | 11 | 4.2 | 4.1 | 3.5 | 2.9 |

**Script of the non-linear least squares (nls) approach modeling following the double component exponential equation:**

```
rm(list=ls())
setwd("~/Folder_with_data")
library(nlstools)

Data=read.table("Data.txt",h=T)

plot(Data$time, Data$percent)

## Von Bertalanffy model, with nls and all parameters
mod0=percent~(100-AE)*exp(-kes*time)+AE*exp(-kel*time)
nls0=nls(mod0,data=data_Cd_Chiro,start=list(kes=0.1,AE=20,kel=0.005))

overview(nls0)
plotfit(nls0,smooth=TRUE)

res0=nlsResiduals(nls0)
plot(res0)
test.nlsResiduals(res0)

region0=nlsConfRegions(nls0)
plot(region0,bounds=TRUE)

contour0=nlsContourRSS(nls0,lseq=50)
plot(contour0,col=TRUE,nlev=10)

shapiro.test(residuals(nls0))

AIC(nls0)
```

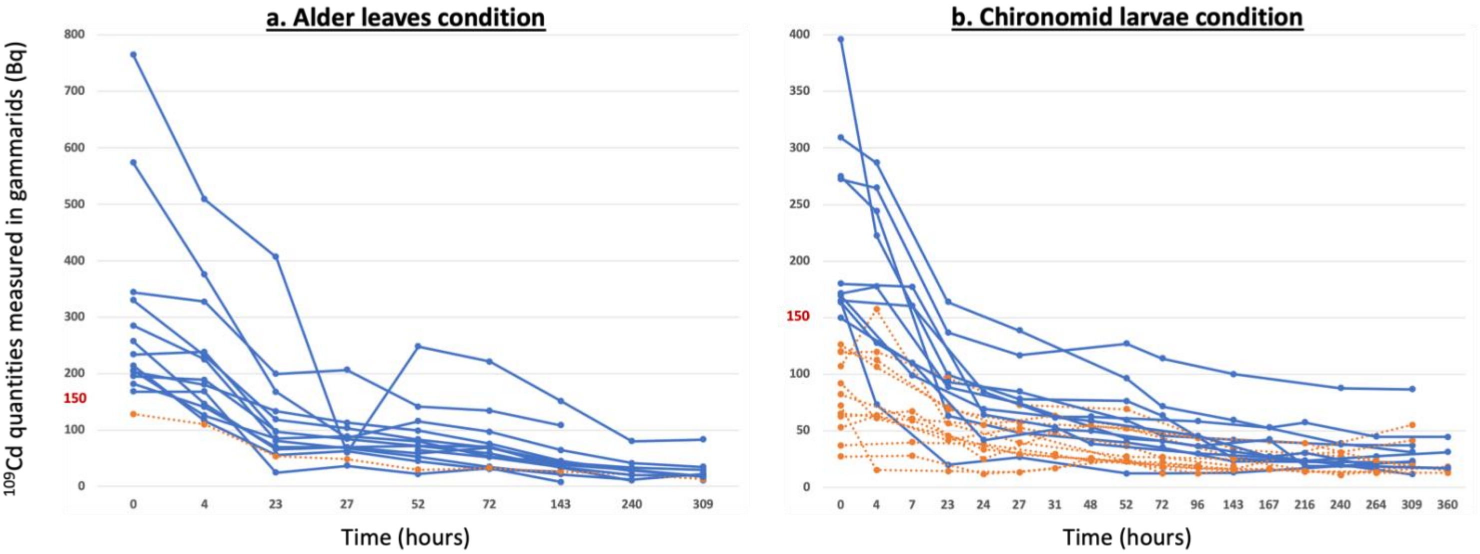


Figure S1. Activity of $^{109}Cd$ measured (Bq) in function of time (hours), from the end of the pulse-chase-feeding (t0) and for a period of approximately 2 weeks. Two types of food were contaminated by $^{109}Cd$, with a) the alder leaves condition and b) the chironomid larvae condition. If, for a gammarid, the activity measured at t0 was less than 150 Bq (orange dot lines) all the data obtained for this gammarid were not used to determine the AE, otherwise (blue lines) the data obtained for this gammarid were retained to estimate the AE.
